# Circulating Cell-Free Chromatin Particles Trigger a Unique Biphasic STING Signaling Program that Drives DNA Damage and Inflammation

**DOI:** 10.64898/2026.07.28.741182

**Authors:** Snehal Shabrish, Swamini Patade, Sushma Shinde, Roohi Yelukar, Gorantla V. Raghuram, Relestina Lopes, Naveen Kumar Khare, Indraneel Mittra

## Abstract

Cell death, DNA damage, and inflammation are closely interconnected processes implicated in ageing, cancer, and inflammatory disorders, yet the endogenous mechanisms linking them remain unclear. We previously identified cell-free chromatin particles (cfChPs), released from dying cells, as biologically active entities that enter neighboring cells and induce DNA damage and inflammation. Here, we show that serum-derived circulating cfChPs are rapidly internalized by human peripheral blood mononuclear cells and trigger a previously unrecognized biphasic STING signaling response. An early phase involves rapid STING trafficking to the perinuclear region and nucleus, with activation of IRF3 and NF-κB preceding detectable DNA damage. This is followed by a later phase characterized by STING phosphorylation, puncta formation, persistent DNA damage, and robust inflammatory cytokine production. Pharmacological inhibition or genetic deletion of STING markedly attenuated these responses. These findings identify extracellular cfChPs as endogenous DNA-damaging agents and reveal biphasic STING signaling as a mechanistic link between cell death, DNA damage, and sterile inflammation.

**One-sentence summary:** Circulating cell-free chromatin particles released from dying cells trigger a previously unrecognized biphasic STING signaling program that mechanistically links cell death to DNA damage and sterile inflammation, providing a potential basis for aging, cancer, and inflammatory diseases.

## Introduction

Cell death, DNA damage and inflammation are closely related biological processes that collectively regulate immune surveillance and disease pathogenesis^1^. While it is known that dying cells release danger-associated molecular patterns (DAMPs), which can trigger sterile inflammatory responses, the endogenous mediators that mechanistically link cell death to DNA damage and innate immune activation remain poorly understood^2,3^. Understanding this relationship has profound biological and clinical implications, as persistent activation of DNA damage and inflammatory pathways contributes to chronic inflammatory diseases, autoimmunity, ageing, and cancer^4^.

Emerging evidence shows that the STING pathway is a central regulator of innate immunity, which detects cytosolic DNA and triggers inflammatory signaling by activating NF-κB and interferon regulatory factors^5–7^. According to current models, genomic instability and DNA damage generate micronuclei or cytosolic DNA fragments that initiate STING signaling, connecting genomic instability and inflammation^5,7,8^. More recently, STING has also been linked to nuclear DNA damage responses, indicating a cyclic interaction in which DNA damage activates STING, and prolonged STING signaling further aggravates genomic instability and inflammatory amplification^6,9^. Despite these advances, the endogenous signal generated during physiological cell death that initiates this signaling cascade has remained incompletely defined^10,11^.

We previously identified cell-free chromatin particles (cfChPs), released from dying cells, as biologically active agents that can be horizontally transferred into healthy cells and accumulate within their nuclei^12–14^. We also reported that this cellular uptake occurs via the phagocytic pathway^13^, and that the internalised cfChPs induce DNA damage and activate inflammatory signaling, suggesting they function as endogenous danger signals that connect cell death to innate immune activation^12,13^. However, the molecular mechanism by which cfChPs couple DNA damage to inflammatory signaling has remained undetermined^15^.

Here, we investigated whether STING signaling might mediate the coordinated DNA damage and inflammatory signaling triggered by cfChPs following their cellular uptake.

The results show that cfChPs trigger a previously unrecognized biphasic STING signaling program that links cell death to DNA damage and sterile inflammation, providing a potential mechanistic basis for ageing, cancer, and inflammatory diseases.

## Materials and Methods

### Ethics statement

The study was approved by the Institutional Ethics Committee of the Advanced Centre for Treatment, Research and Education in Cancer (ACTREC), Tata Memorial Centre, Mumbai, India (Approval No. 900990). Peripheral blood samples were obtained from healthy adult volunteers after written informed consent in accordance with institutional guidelines and the Declaration of Helsinki.

### Isolation of peripheral blood mononuclear cells

Peripheral blood was collected from healthy volunteers with no history of febrile or significant illness during the preceding three months. PBMCs were isolated from heparinized blood by Ficoll-Hypaque density-gradient centrifugation using the standard procedure.

### Cell Culture

Isolated PBMCs were cultured in Dulbecco’s Modified Eagle Medium (DMEM) supplemented with 10% fetal bovine serum at 37°C in a humidified atmosphere containing 5% CO₂. In some experiments, THP-1 wild-type (WT) and STING-knockout (KO) cells (InvivoGen) were used and cultured in RPMI-1640 supplemented with 10% fetal bovine serum and 50 μM β-mercaptoethanol and maintained as above. Cell lines were routinely tested and confirmed to be free of mycoplasma contamination and were authenticated by short tandem repeat profiling.

For estimation of inflammatory cytokines, THP-1 cells were differentiated into macrophages using 100 ng/ml phorbol-12-myristate-13-acetate (TPA) for 24 h, and cultured for 48 h without TPA.

### Isolation of cell-free chromatin particles from serum

For isolation of cell-free chromatin particles (cfChPs), serum samples from healthy volunteers were subjected to ultracentrifugation at 694,000 × g at 4°C for 16 hours. Chromatin from the pellets were extracted using the ChromaFlash™ Chromatin Extraction Kit as per the manufacturer’s instructions. Briefly, the pellet was lysed in a lysis buffer supplemented with a protease inhibitor cocktail and incubated on ice for 10 minutes, followed by ultracentrifugation. Extraction buffer supplemented with a protease inhibitor cocktail was added to the pellet, followed by sonication and clarification by centrifugation. Purified cfChPs were aliquoted and stored at −80°C until use. A nucleosome-specific sandwich ELISA kit (Cell Death Detection ELISA^PLUS^ kit, Roche Diagnostics GmbH, Germany) was used to confirm the presence of cfChPs. A representative electron microscopy image of the cfChPs is given in **Supplementary Figure 1**. The isolated cfChPs were quantified by DNA content using a PicoGreen quantification assay.

### Fluorescent Labeling of cfChPs

For cellular uptake studies, cfChPs were fluorescently dually labelled in their DNA and histone H4 components as previously described by us^12^. DNA was labeled with Platinum Bright™ 550 Red Nucleic Acid Labelling Kit (Kreatech Diagnostics, Cat # GLK-004) and histone H4 with ATTO 488 NHS-ester (ATTO-TEC GmbH, Cat # AD488-35). PBMCs were treated with 400 pg of dually labeled cfChPs, washed with PBS, and examined using an Applied Spectral Imaging platform.

### Cell Treatments

PBMCs (1 × 10⁶ cells) and THP-1 cells (0.5 × 10⁶ cells) were seeded in 24-well plates and allowed to recover overnight. Cells were treated with serum-derived unlabelled cfChPs equivalent to 400 pg of DNA. To evaluate the role of STING signaling, PBMCs were either pre-treated with the STING inhibitor H-151, or the wild-type or STING-deficient THP-1 cells were used.

### Quantitative Real-Time PCR

Total RNA was isolated using silica membrane-based purification, and cDNA was synthesised from 1 μg RNA using standard procedures. Gene expression analyses were performed using SYBR Green-based quantitative PCR on a QuantStudio 12K Flex system. Relative transcript abundance was calculated using the 2−ΔΔCt method following normalisation to the corresponding housekeeping gene.

### Flow Cytometry

Flow cytometry was performed to assess immune cell activation, DNA damage responses, STING activation and inflammatory markers.

#### Estimation of markers of T-cell activation

In a time-course analysis, cells were surface-stained with fluorochrome-conjugated antibodies against lineage (CD45, CD3) and activation markers (CD69 and HLA-DR), (Commercial sources and catalogue numbers of the antibodies are given in **Supplementary Table 1**) and acquired using Attune NxT (ThermoFisher Scientific, USA) flow cytometer. T cells were identified as CD3⁺ lymphocytes, and expression levels were quantified as percentage-positive cells using flow cytometry. Data were analysed using FlowJo software (Version 10; FlowJo LLC, Ashland, OR, USA)

#### Estimation of DNA damage response, inflammatory markers, and pSTING expression

Peak time-points of mRNA expression were determined by RT-PCR in time course analysis. PBMCs were fixed and permeabilized with either 4% formaldehyde and 90% methanol or cytofix / cytoperm kit (Becton Dickinson) based on the manufacturer’s instructions. Cells were stained using antibodies against γ-H2AX, granzyme B, JUN-B, c-JUN, FOS-B, and pSTING (Commercial sources and catalogue numbers of the antibodies are given in **Supplementary table 1**). Cells were stained with appropriate secondary antibodies **(Supplementary Table 1)**. Cells were acquired on the Attune NxT flow cytometer (ThermoFisher Scientific, USA). Data were analysed using FlowJo software (Version 10; FlowJo LLC, Ashland, OR, USA)

#### Cytokine and Chemokine Analysis

Cell-culture supernatants were collected 48 h after treatment and stored at −80°C until analysis. A LEGENDplex™ Multi-Analyte Flow assay kit (BioLegend, San Diego, CA, USA) was used as per the manufacturer’s instructions to quantify levels of secreted cytokines in culture supernatants. At least 4,000 cytometric beads were acquired on Attune NxT flow cytometer (ThermoFisher Scientific, USA) and analysed using LEGENDplex Data Analysis Software (BioLegend, San Diego, CA, USA).

### Immunofluorescence

PBMCs were treated with cfChPs (400 pg) and harvested at appropriate time points. Cells were cytospun onto glass slides, fixed with paraformaldehyde and stained using primary antibody against γ-H2AX, STING, pSTING, pNF-κB and pIRF3 and appropriate secondary antibodies by the indirect immunofluorescence method as described by us earlier^16^. The commercial sources of these antibodies are given in **Supplementary Table 1**. Slides were mounted in VectaShield DAPI (Vector Labs, USA). Images were acquired using an Applied Spectral Imaging platform, and fluorescence intensity was quantified using Fiji/ImageJ software. Approximately 100-200 cells were analysed in each case.

### Confocal microscopy

For visualising the trafficking of STING protein in response to cfChPs treatment, slides were acquired using confocal microscopy. Images were acquired using a 63x oil objective on the Nikon AX Confocal Microscope System (Nikon Corporation, Japan).

### Statistical Analysis

Data are presented as mean ± sem. Statistical analyses were performed using GraphPad Prism 8. Comparisons between two groups were conducted using two-tailed unpaired Student’s t-tests unless otherwise indicated in the figure legends. Differences were considered statistically significant at P < 0.05.

## Results

### Serum-derived cfChPs are rapidly internalised by PBMCs

Treatment of PBMCs with dual-fluorescent labelled cfChPs resulted in their rapid uptake by 2 h and nuclear accumulation by 4 h as visualised by immunofluorescence **(Supplementary Figure 2)**.

### cfChPs induce biphasic activation of STING signaling

We next investigated whether cfChPs uptake by PBMCs activates the STING pathway, a central regulator of cellular responses to the presence of aberrant intracellular chromatin. Temporal analysis of STING activation in PBMCs following exposure to cfChPs by immunofluorescence analysis revealed rapid redistribution of intracellular STING. STING accumulated around the nucleus in the perinuclear region within 1 h of cfChPs treatment, followed at 2 h by its translocation to the nucleus (**Figure 1a).** This early phase of STING activation (2 h) was accompanied by nuclear translocation of phosphorylated IRF3 (pIRF3) and phosphorylated NF-κB (pNF-κB), indicating that the initial phase of STING signaling pathways were activated **(Supplementary Figure 3)**. A second phase of STING activation became evident by 12 h, characterized by redistribution of STING as discrete cytoplasmic puncta, a hallmark of prolonged STING signaling activation **(Figure 1a)**. This was associated with a marked increase in expression of pSTING at 12 h and 24 h as determined by flow cytometry in a time-course analysis **(Figure 1b).** These results were validated by immunofluorescence, showing the phosphorylation of STING after 24 h exposure to cfChPs **(Figure 1c).**

**Figure 1.**
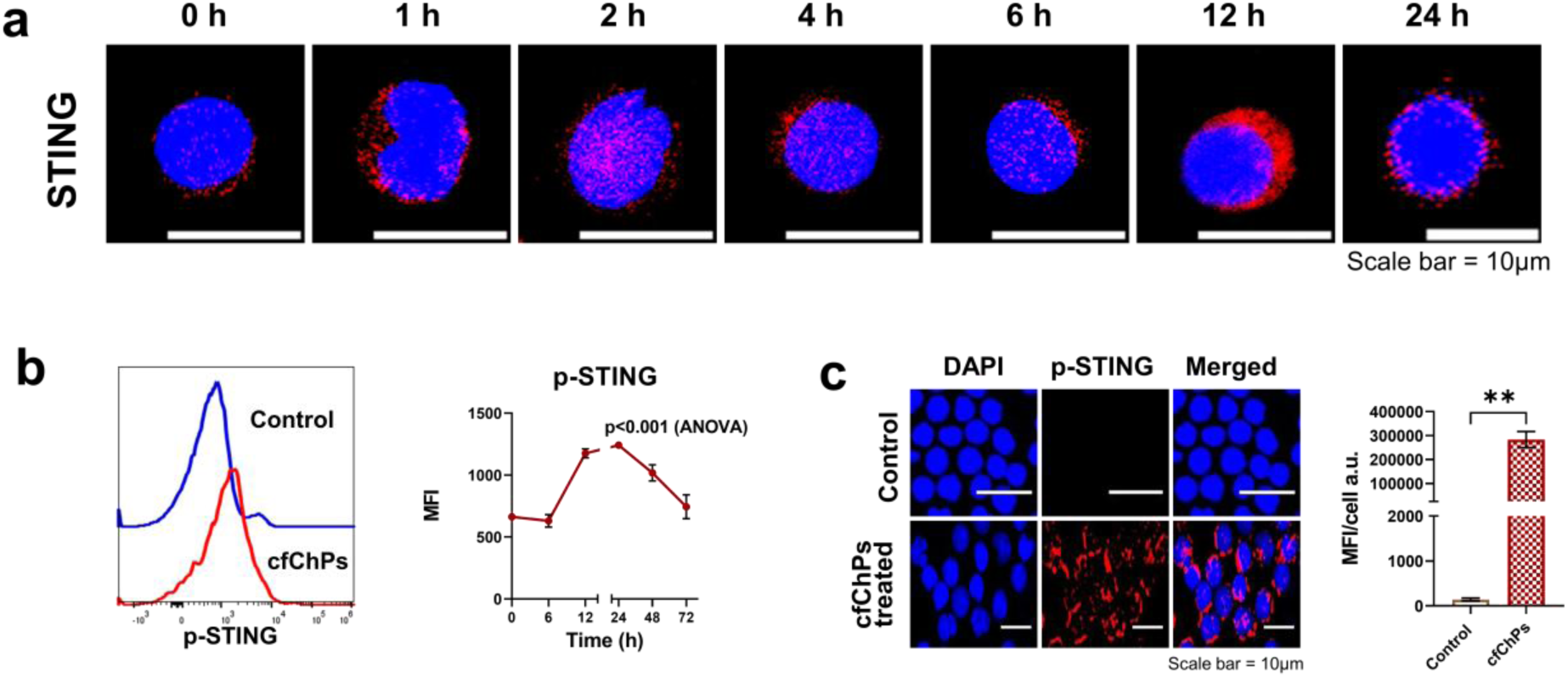
Serum-derived cfChPs induce biphasic activation of STING in human PBMCs. **(a)** Representative confocal immunofluorescence images showing the temporal redistribution of STING (red) in human PBMCs following treatment with serum-derived cfChPs. STING localized predominantly in the cytoplasm of untreated cells, rapidly redistributed to the perinuclear region within 1 h of cfChPs exposure, followed by nuclear accumulation at 2 h. By 12 h, STING formed discrete cytoplasmic puncta, indicative of sustained STING activation. **(b)** Flow cytometric analysis of pSTING expression comparing untreated control cells (blue) and cfChP-treated cells (red) shows increased pSTING expression following treatment. The line graph represents the temporal quantification of MFI of pSTING expression in a time-course analysis **(c)** Representative immunofluorescence images of pSTING (red) in untreated and cfChP-treated PBMCs show increased pSTING expression following treatment. The histogram shows the relative upregulation of pSTING in cfChP-treated cells compared to the untreated control. Data are presented as mean ± SEM. Statistical significance was determined using one-way ANOVA and two-tailed unpaired Student’s t-test in GraphPad Prism 8. **\*\*** p<0.01

### cfChPs induce DNA damage

In confirmation of our earlier reports based on mouse fibroblast cells^12,13^, treatment of PBMCs with serum-derived circulating cfChPs elicited a robust DNA damage response, as determined by phosphorylation of H2AX, a sensitive marker of DNA double-strand breaks. Time-course analyses by immunofluorescence microscopy and flow cytometry revealed a marked increase in γ-H2AX-positive cells as early as 2 h, with signal intensity increasing progressively over time up to 24 h (ANOVA, P < 0.0001). At a latter time-point, the treated cells exhibited a >20-fold increase in γ-H2AX-positive cells **(Figure 2a, Supplementary Figure 4)**.

**Figure 2.**
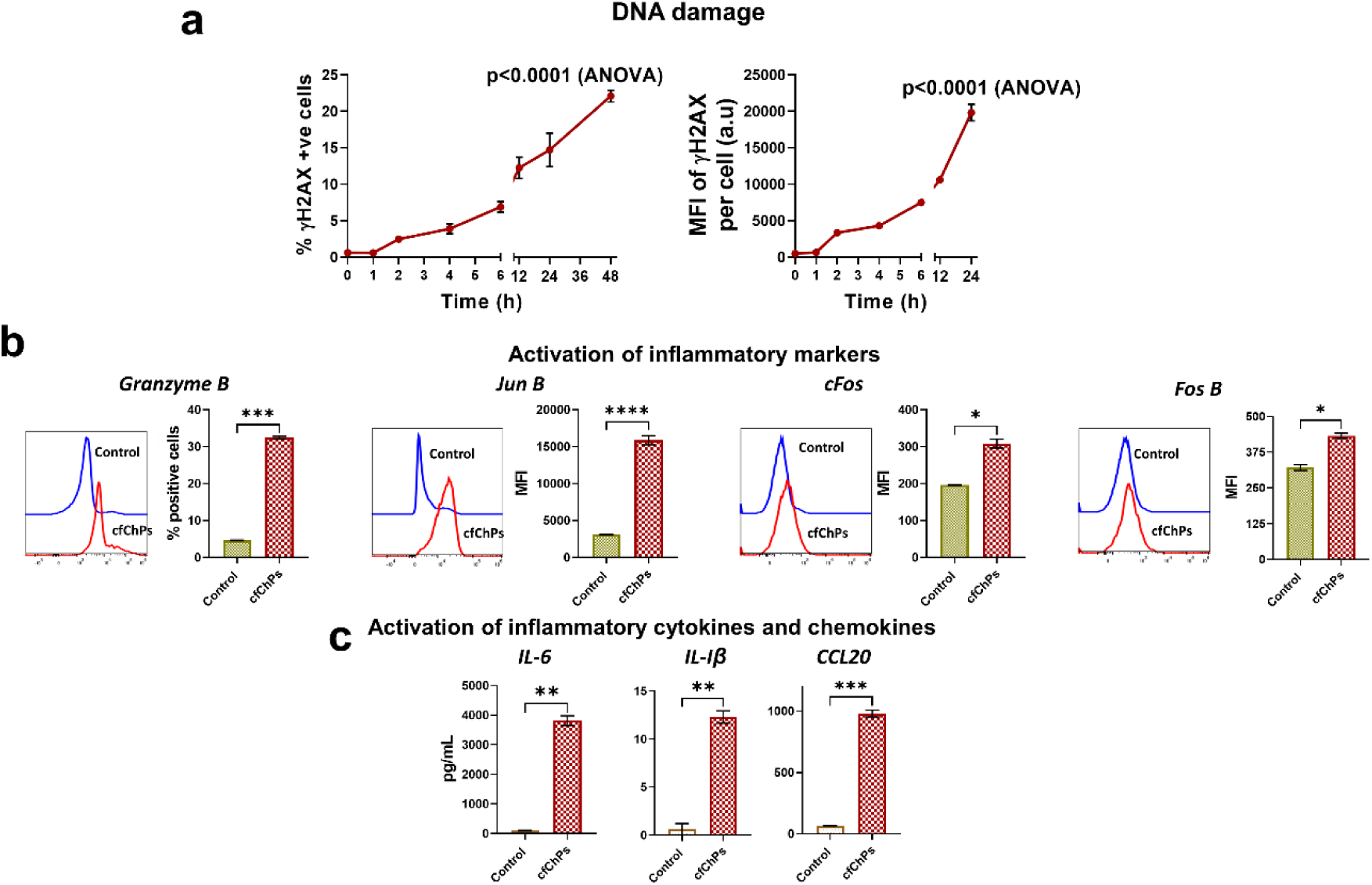
Serum-derived cfChPs induce DNA damage and activate an immune response. **(a)** Line graphs represent the temporal increase in the expression of γ-H2AX in a time-course experiment using flow cytometry (left-hand side) and immunofluorescence (right-hand side). **(b)** Upregulation of inflammatory marker expression in PBMCs treated with cfChPs. Samples were analyzed by flow cytometry, and respective protein expression was analyzed on untreated (blue line) and cfChPs treated (red line) PBMCs. Histograms represent quantitative analysis (MFI) of respective inflammatory markers in PBMCs treated with cfChPs. **(c)** Histograms show that the production of cytokines by PBMCs in response to cfChPs. Data were analyzed using two-tailed unpaired Student’s *t*-tests in GraphPad Prism 8. **\*** p<0.05, **\*\*** p<0.01, **\*\*\*** p<0.005, **\*\*\*\*** p<0.0001.

### cfChPs activate an immune response

We next investigated the relation between DNA damage and inflammation. We examined T-cell activation, expression of inflammatory markers, and secretion of inflammatory mediators following treatment of PBMCs with cfChPs. Flow cytometric analysis showed that cfChPs treatment significantly increased the expression of the activation markers CD69 and HLA-DR on T cells, with the maximal expression occurring at 24 h (P < 0.0001) (**Supplementary Figure 5a**). Consistent with T-cell activation, cfChPs also induced the expression of multiple inflammatory genes, including Granzyme B, JunB, FosB, and c-Fos (**Supplementary Figure 5b**). Granzyme B mRNA expression peaked at 24 h, whereas JunB, FosB, and c-Fos reached maximal expression at 48 h (**Supplementary Figure 5b**). Increased protein expression of all four markers at their respective peak time points was confirmed by flow cytometry (**Figure 2b**). Treatment with cfChPs resulted in a significant increase in the proportion of granzyme B-positive cells compared with untreated controls (P< 0.001) **(Figure 2b),** and also induced a marked increase in expression of inflammatory signaling markers, namely, JunB (P< 0.0001), c-Fos (P< 0.05) and FosB (P< 0.05), as demonstrated by a pronounced increase in mean fluorescence intensity (MFI) compared to untreated PBMCs.

Additionally, cfChPs treatment resulted in a significant increase in the secretion of the pro-inflammatory cytokines IL-1β (P < 0.01) and IL-6 (P < 0.01), together with the chemokine CCL20 (P < 0.001) (**Figure 2c**). Collectively, these findings demonstrate that serum-derived circulating cfChPs not only induce DNA damage but also activate a robust inflammatory response in human PBMCs.

### STING signaling is critical for cfChP-induced DNA damage and inflammatory response

Given that cytosolic DNA is a potent activator of the STING pathway, and serum-derived cfChPs can induce DNA damage and inflammatory signaling, we investigated whether these activities of cfChPs are STING-mediated responses. We employed two approaches: pharmacological inhibition of STING using H151 in human PBMCs, and THP-1 cells which are genetically ablated for STING. We investigated whether STING inhibition would abrogate inflammation accompanying DNA damage.

Pharmacological inhibition of STING markedly attenuated the inflammatory response induced by cfChPs. PBMCs treated with cfChPs in the presence of H151 showed significantly reduced production of the pro-inflammatory cytokines IL-1β (*P* < 0.05) and IL-6 (*P* < 0.05), as well as the chemokine CCL20 (*P* < 0.01), compared with cells treated with cfChPs in the absence of H151. These responses were determined by flow cytometry-based cytometric bead array **(Figure 3a)**.

**Figure 3.**
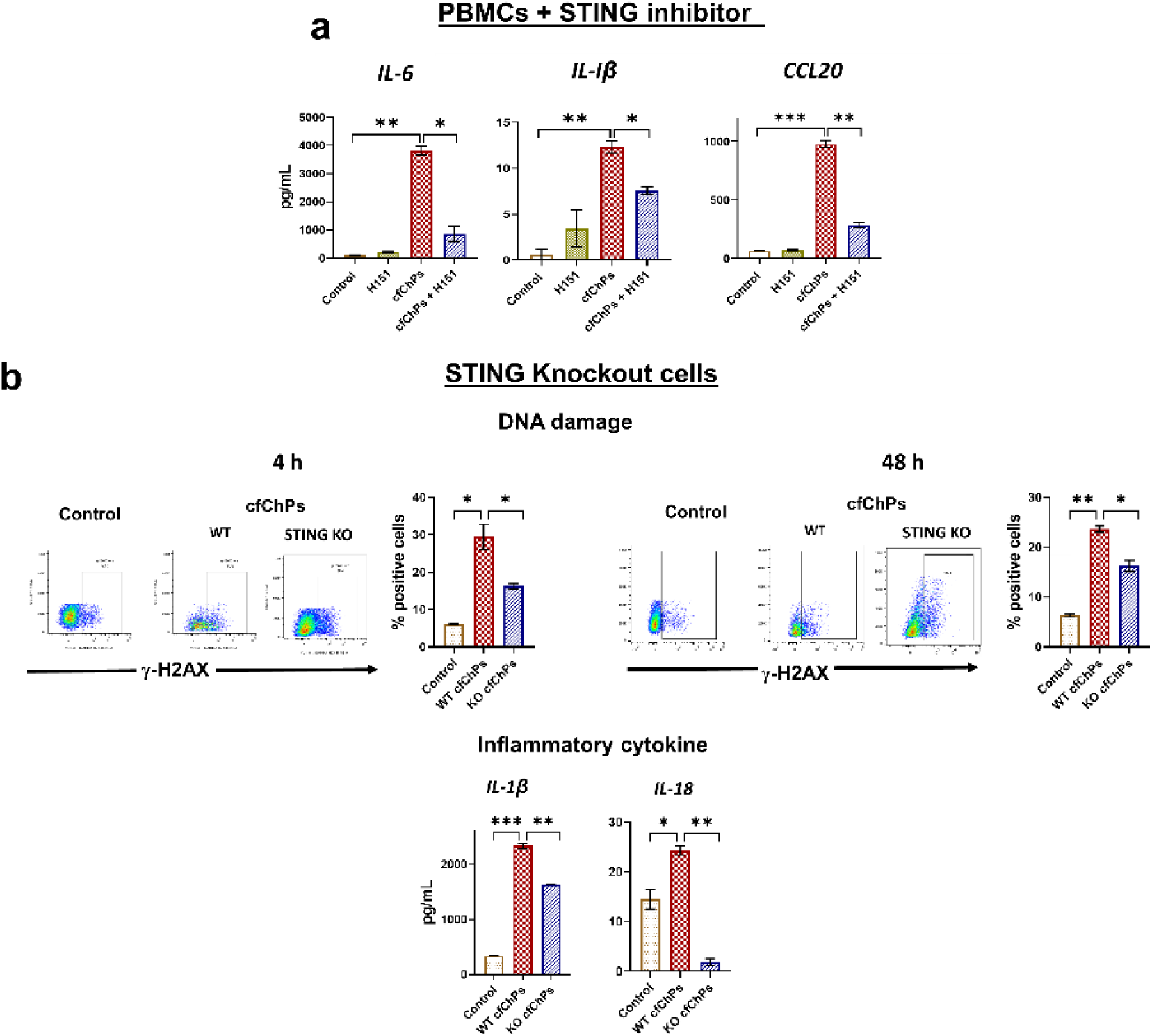
STING is required for cfChP-induced DNA damage and inflammatory responses. **(a)** Pharmacological inhibition of STING with H151 attenuates cfChP-induced cytokine production in PBMCs. Histograms show that H151 significantly reduced cfChP-induced secretion of IL-6, IL-1β, and CCL20. **(b)** Genetic ablation of STING in the THP-1 cell line suppresses cfChP-induced DNA damage (upper panel) and inflammatory cytokine secretion (lower panel). Representative flow cytometry plots (left-hand side) and quantification (right-hand side) of γ-H2AX-positive cells at 4 h and 48 h following cfChPs treatment in wild-type (WT) THP-1 cells and STING-knockout (STING-KO) THP-1 cells. The lower panel shows histograms indicating reduced levels of IL-1β and IL-18 in cfChP-treated STING-knockout THP-1 cells compared to wild-type cells. Data are presented as mean ± SEM. Statistical significance was determined by two-tailed unpaired Student’s *t*-tests in GraphPad Prism 8. **\*** p<0.05, **\*\*** p<0.01, **\*\*\*** p<0.005.

To further establish the role of the STING protein in cfChPs-mediated DNA damage and inflammatory response, we compared the effects of cfChPs in wild-type and STING-knockout THP-1 cells. Loss of STING significantly reduced the DNA damage response, indicated by markedly reduced γ-H2AX positive cells detected at both 4 h and 48 h following cfChPs exposure (*P* < 0.05) **(Figure 3b).** Consistent with these findings, a significant reduction in levels of the pro-inflammatory cytokines IL-1β and IL-18 was observed in the STING KO THP-1 cells compared with the wild-type cells following cfChPs treatment (*P* < 0.01) **(Figure 3b)**. Collectively, these results demonstrate that STING is a critical mediator of both DNA damage response and inflammatory signaling induced by circulating cfChPs.

## Discussion

Ageing, cancer, and many inflammatory disorders are driven by the closely interconnected biological processes of cell death, DNA damage, and inflammation^1^. Although these processes have long been recognised to influence one another, the endogenous mechanisms that mechanistically link them remain poorly understood^4^. While dying cells release DAMPs capable of initiating sterile inflammatory responses, the endogenous mediator that links physiological cell death to DNA damage and activation of innate immune signaling has remained to be clearly defined^2,3^.

Here, we identify circulating cfChPs released from dying cells as endogenous mediators that link cell death, DNA damage, and inflammation through a previously unrecognised pattern of STING activation. Whereas the prevailing model places STING activation downstream of DNA damage^6–9^, our findings support a fundamentally different sequence wherein extracellular cfChPs initiate STING activation before the onset of a robust DNA damage response. We demonstrate that cfChPs are rapidly internalised by recipient immune cells through horizontal transfer, triggering an early phase of STING activation that precedes detectable DNA damage response. This is subsequently accompanied by activation of the downstream transcription factors IRF-3 and NFκB. These early events are followed by a second phase of STING activation characterised by STING phosphorylation and amplification of inflammatory signaling. These observations place cfChPs at the apex of this signaling cascade and identify them as physiological danger signals capable of co-ordinating multiple stress responses.

Our findings extend previous observations that circulating cfChPs released during physiological cell death are biologically active entities rather than inert by-products of tissue injury. We show that cfChPs generated from the billions of cells that die daily and circulate in human blood are efficiently internalised by human PBMCs and induce DNA double-strand breaks (DSBs), immune activation, and the production of inflammatory cytokines and chemokines. These observations are consistent with our earlier studies showing the rapid horizontal transfer of cfChPs into recipient cells and their ability to induce DNA damage^12,14,17^. They also support the emerging concept that endogenous DNA double-strand breaks are potent initiators of sterile inflammation^15^. Collectively, these findings identify circulating cfChPs as a physiologically important endogenous source of DNA damage that can initiate and sustain inflammatory signaling.

The most significant finding of our study is the identification, to our knowledge for the first time, of a biphasic pattern of STING activation induced by cfChPs. Within one hour of cfChP exposure, STING rapidly redistributed to the nucleus and perinuclear region, subsequently accompanied by nuclear translocation of phosphorylated NF-κB and IRF3, before any detectable evidence of DNA damage. This early phase was followed by a second phase characterised by the formation of perinuclear STING puncta and phosphorylation of STING, coinciding with robust cytokine production and immune activation. The temporal separation of these events suggests that STING performs distinct functions during the two phases of the response. The early STING trafficking may facilitate the initiation of DNA damage response and transcriptional reprogramming, whereas subsequent puncta formation and phosphorylation sustain inflammatory signaling. These findings are consistent with growing evidence that STING possesses non-canonical nuclear functions in genome maintenance and the DNA damage response, in addition to its well-established role as a cytosolic adaptor of innate immune signaling^18,19^.

The mechanism by which internalised cfChPs activate STING before evidence of DNA damage remains to be elucidated. One possibility is that endocytosed cfChPs while transiently passing through the cytosol before nuclear accumulation, result in activation of the canonical cGAS-STING pathway. The other possibility is the hypothesis that STING itself can localise to the inner nuclear membrane and directly activate an inflammatory response^9,18^. Our observation of rapid nuclear accumulation of cfChPs suggests they may provide the necessary trigger for the perinuclear redistribution of STING, as suggested by the above hypothesis. Future studies investigating the contributions of nuclear STING complexes, cGAS, and DNA repair proteins will be important for defining this pathway.

The present study has certain limitations. Although primary human PBMCs and STING-deficient THP-1 cells provide compelling evidence for a STING-dependent mechanism, the precise molecular intermediates connecting cfChPs to STING activation remain to be determined. Furthermore, our findings were obtained predominantly in ex vivo cell systems, and validation in relevant animal models will be required to establish the physiological significance of this pathway to tissue injury, ageing, and chronic inflammatory disease.

Taken together, our findings suggest a revised model in which circulating cfChPs released during physiological cell death function as endogenous danger signals that initiate a biphasic programme of STING signaling, leading first to DNA damage and subsequently to amplification of inflammatory signaling. This work potentially identifies the missing biological link between cell death, DNA damage, and sterile inflammation, providing a conceptual framework for understanding how physiological cell turnover contributes to chronic inflammatory diseases. Given the central role of cfChPs in this pathway, therapeutic strategies that neutralise extracellular chromatin may offer new opportunities for treating inflammatory disorders, autoimmunity, ageing, and cancer. Indeed, our previous studies demonstrating the therapeutic efficacy of extracellular chromatin inactivation in experimental models of sepsis, ageing^20–23^, and cancer provide proof-of-concept for targeting this pathway.

Given that several hundred billion cells die in the body every day^24,25^, our results suggest that circulating cell-free chromatin particles released from these dying cells are the natural triggers for the DNA damage and inflammation which are mediated by a unique biphasic STING mechanism, providing a potential basis for aging, cancer, and inflammatory diseases.

## Acknowledgements

The authors thank the personnel in the flow cytometry and digital imaging facility at ACTREC-TMC for their technical support.

## Funding

The authors gratefully acknowledge the financial support provided by the Anusandhan National Research Foundation (ANRF), Government of India (formerly Science and Engineering Research Board (SERB)) under the Core Research Grant scheme (CRG/2023/004470).

## Author contributions

S.Sh.: Conceptualization, Methodology, Investigation, Data Curation, Funding Acquisition, Writing – Original Draft. S.P.: Investigation, Formal Analysis. S.Shi.: Investigation (Immunofluorescence). R.Y.: Investigation (qRT-PCR). G.V.R.: Supervision (Microscopy). R.L.: Investigation (Tissue culture). N.K.K.: Investigation (Chromatin isolation). I.M.: Conceptualization, Methodology, Supervision, Project Administration, Writing – Review & Editing.

## Competing interests

The authors declare that they have no competing interests.

## Data and materials availability

All data supporting the findings of this study are available within the article and supplementary materials.

## AI Use Statement

ChatGPT (OpenAI) was used only to assist with language editing, improving readability, and organising the manuscript. All scientific ideas, interpretations, literature selection, and final content are the responsibility of the author.

## Notes

### Competing Interest Statement

The authors have declared no competing interest.

