## Supplementary figures for "Circulating Cell-Free Chromatin Particles Trigger a Unique Biphasic STING Signaling Program that Drives DNA Damage and Inflammation"

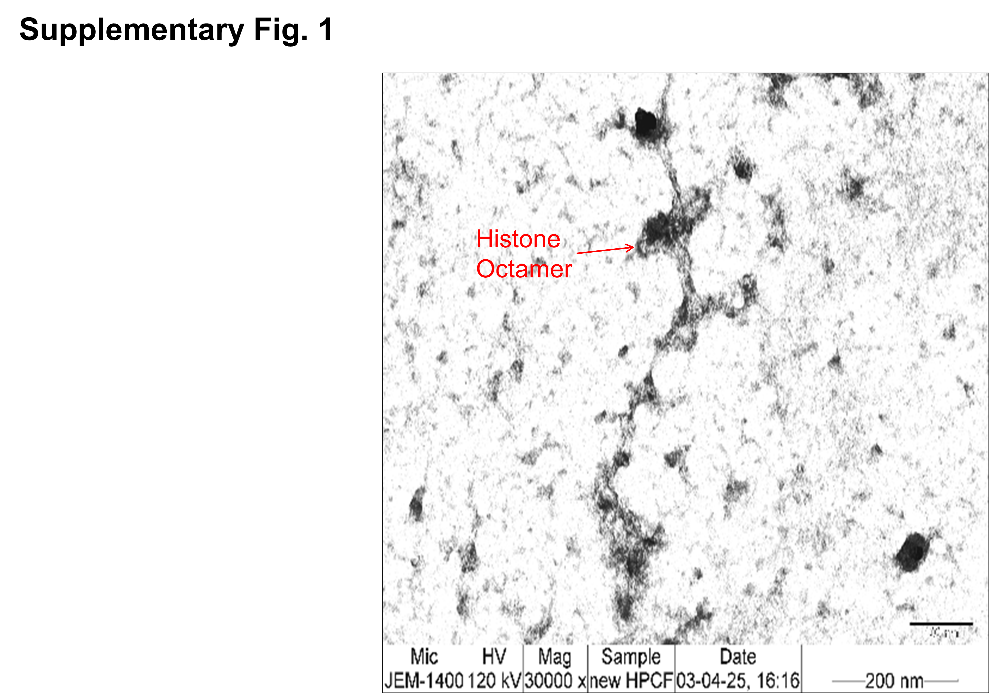

**Supp. Fig. 1.** A representative EM image of cfChPs isolated from human serum showing beads-on-a-string appearance typical of chromatin.

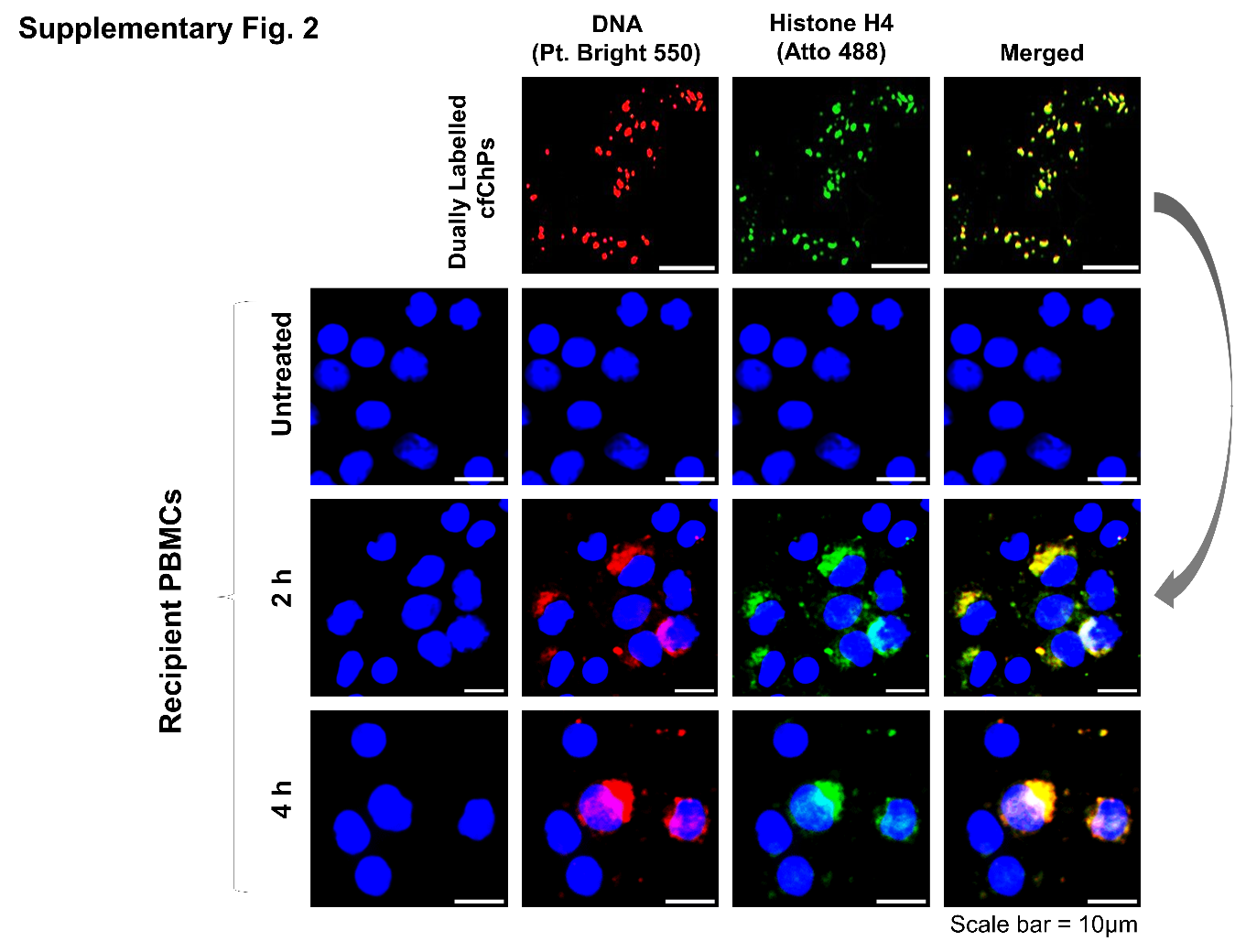
**Supp. Fig. 2. Serum-derived cfChPs are rapidly internalised by PBMCs.** Serum-derived cfChPs were dually labelled in their DNA and histones with Platinum Bright 550 (red) and ATTO-TEC-488 (green), respectively. PBMCs were treated with the dually labelled cfChPs (400 pg), and fluorescent microscopy images show rapid uptake by 2 h and their nuclear accumulation by 4 h.

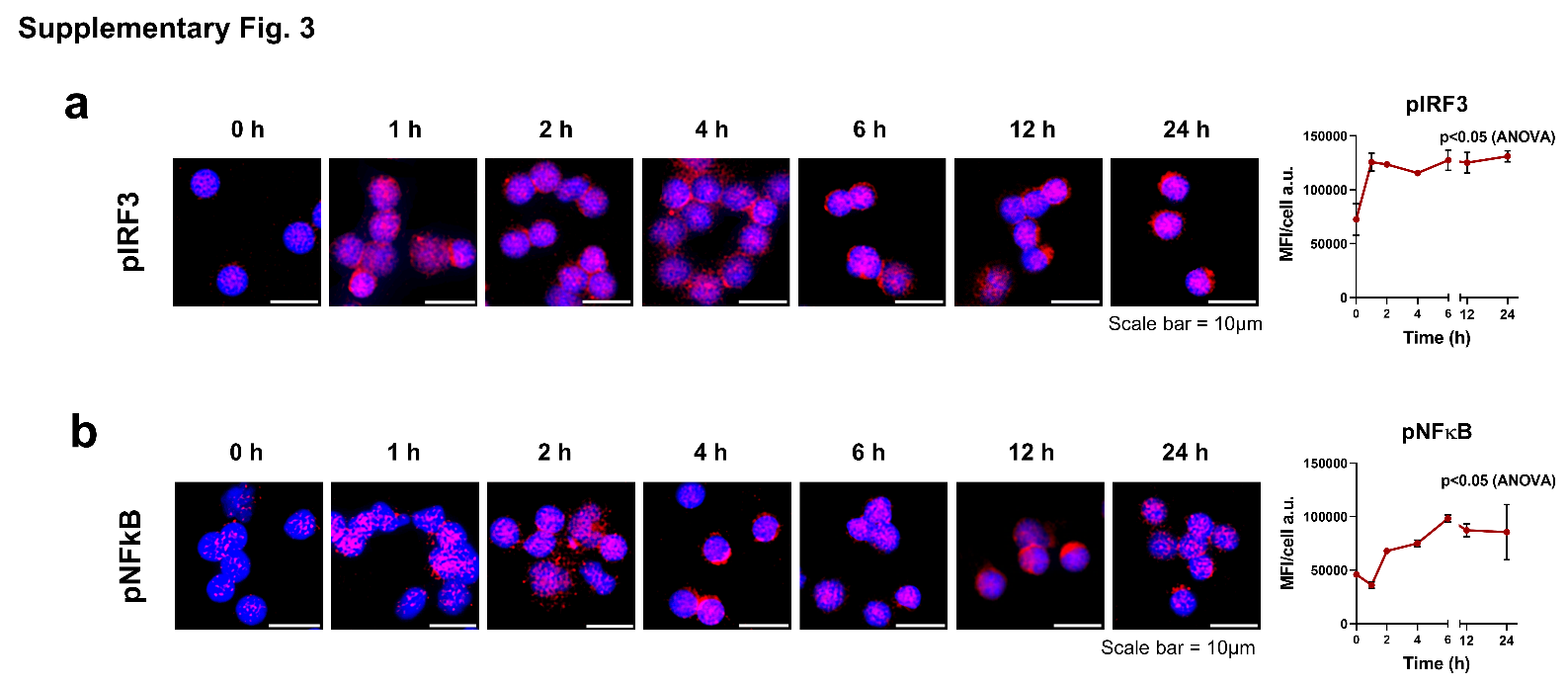

**Supp. Fig. 3. Serum-derived cfChPs activate the STING signaling pathway.** Representative immunofluorescence images showing nuclear translocation of **(a)** pNFκb and **(b)** pIRF3 in PBMCs treated with cfChPs in a time-course analysis. The respective line graphs represent the temporal increase in MFI of nuclear expression of p-NFκb and pIRF3.

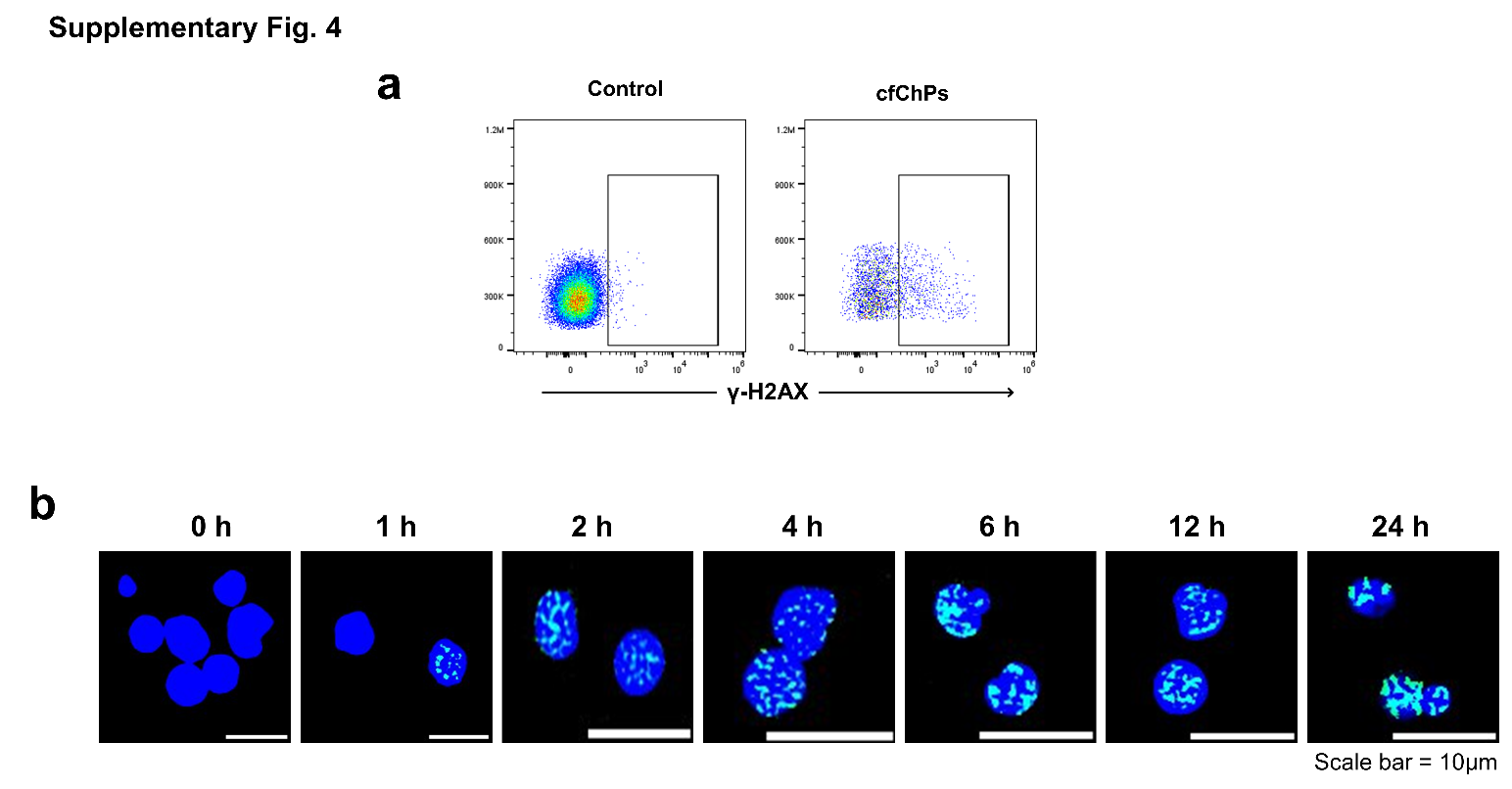

**Supp. Fig. 4. Serum-derived cfChPs induce DNA damage. (a)** Flow cytometry representative images showing upregulation of γ-H2AX after 24 h treatment of PBMCs with cfChPs. **(b)** Immunofluorescence representative images showing temporal progressive upregulation of γ-H2AX in PBMCs treated with cfChPs.

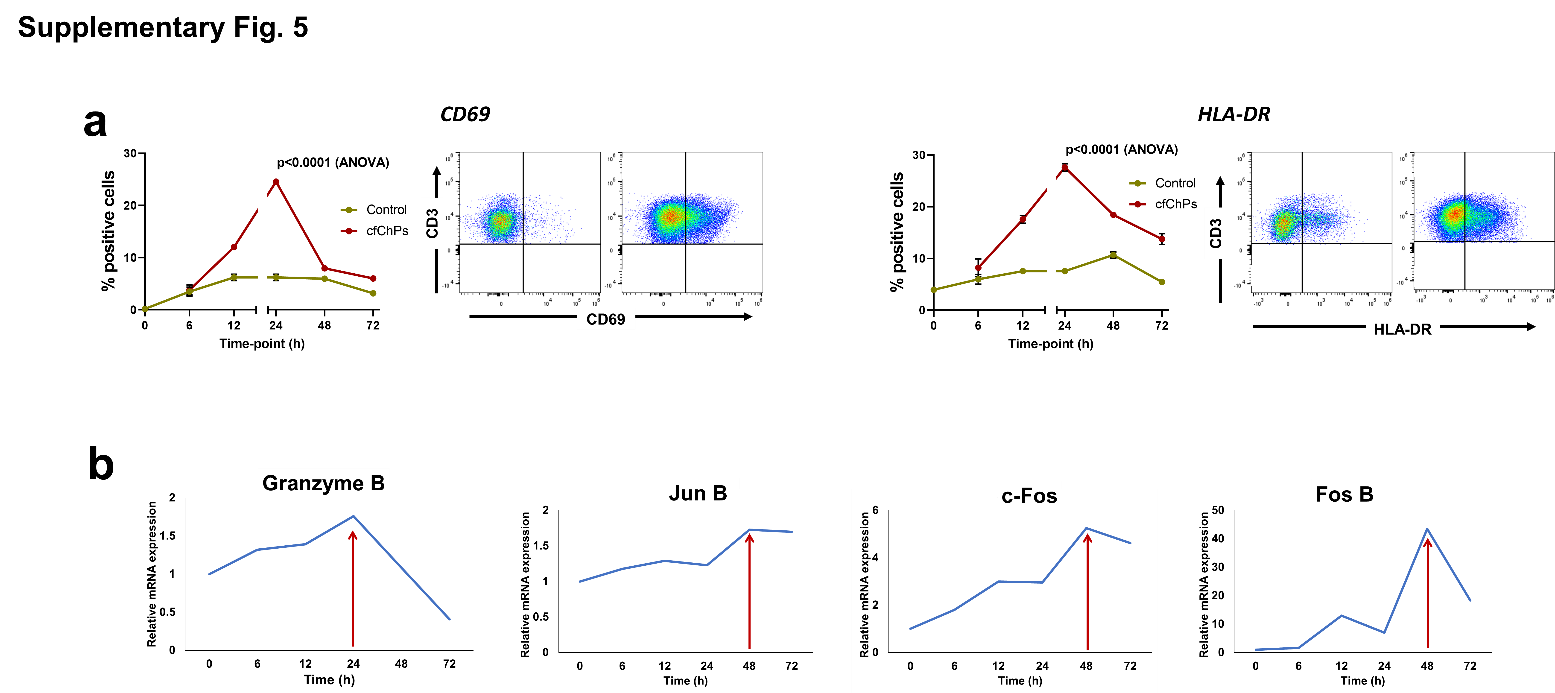

**Supp. Fig. 5. Serum-derived cfChPs activate an immune response. (a)** Line graphs represent the expression of CD69 (left-hand panel) and HLA-DR (right-hand panel) in a time-course experiment as detected by flow cytometry (green line represents untreated cells; red line represents cfChP-treated cells). Representative flow cytometry plots represent the relative upregulation of CD69 expression on T-cells and NK-cells in response to cfChPs at the peak time-point (24 h). **(b)** Upregulation of stress-related markers in human PBMCs following treatment with cfChPs (10 ng). mRNA expression of six stress-related markers in PBMCs was detected by qRT-PCR and analyzed using a comparative C_T_ method. Line graphs represent the results of time course analyses calculated as expression fold [2^(-ΔΔC_T_)].
