## Supplementary table 1 for "Circulating Cell-Free Chromatin Particles Trigger a Unique Biphasic STING Signaling Program that Drives DNA Damage and Inflammation"

| **Reagent** | **Catalogue no.** | **Company** |
| --- | --- | --- |
| CD3 | 563725 | BD Horizon |
| CD45 | 560178 | B.D Pharmingen |
| CD69 | 562884 | BD Horizon |
| HLA-DR | 559866 | B.D Pharmingen |
| γ-H2AX (Flow) | 5763 | Cell Signaling Technology |
| γ-H2AX (IF) | 05-636-1 | EMD Millipore Corp, USA |
| Granzyme B | 571117 | BD Horizon |
| Jun B | NBP3-08391 V | Novus Biologicals |
| c-Fos | 2250S | Cell Signaling Technology |
| FosB | 2251S | Cell Signaling Technology |
| STING | MAB7169 | R& D Systems |
| p-STING | 40818S | Cell Signaling Technology |
| p-NFκb | MA5-15160 | Invitrogen |
| p-IRF3 | 29047 | Cell Signaling Technology |
| LEGENDplex™ Mul­ti-Analyte Flow assay kit – Cytokine Bead Assay | 740809 | BioLegend, San Diego, CA, USA |
| LEGENDplex™ Mul­ti-Analyte Flow assay kit – Chemokine Bead Assay | 740985 | BioLegend, San Diego, CA, USA |
| Goat anti-Mouse IgG (H+L) Secondary Antibody, DyLight 633 | 35512 | Thermo-Fisher Scientific, USA |
| Goat anti-Rabbit IgG (H+L) Secondary Antibody, DyLight 633 | 35562 | Thermo-Fisher Scientific, USA |

**Supplementary table 1.**

**Details of antibodies used in this study (As per vendors’ specifications)**
